# The Fengshu Large Model for Wugu Fengdeng: An Innovation Engine for Knowledge Integration in the Soybean Field

**DOI:** 10.1101/2025.06.30.662271

**Authors:** Maoyang Wang, Juncheng Ling, Pengsheng Qiao, Mengxue Yang, Dongbo Liu, Xiaodong He, Zhenbang Hu, Rongsheng Zhu, Qingshan Chen

## Abstract

Against the backdrop of global population growth and the continuous escalation of food demand, the acceleration of agricultural modernization has emerged as the core pathway to safeguard food security. As the world’s fourth-largest food crop, soybean (Glycine max) possesses multiple strategic values for food security, livestock feed, and industrial raw materials, thanks to its high protein content (accounting for over 40% of the dry weight of seeds) and oil resource attributes. However, the soybean industry is confronted with multiple challenges: the long cycle of genetic breeding (8-10 years required by traditional methods), annual losses from diseases and pests reaching 40% (data from FAO 2023), and the reliance on empirical decision-making in field management, all of which urgently call for intelligent solutions.At present, agricultural knowledge is experiencing explosive growth - more than 4,000 new soybean-related literatures are added annually in PubMed, and agricultural technology Q&A platforms (Zhihu, Baidu Tieba, etc.) generate over 3,000 daily questions. However, this multi-source heterogeneous knowledge is scattered in books, literatures, Q&A communities, and gene databases, lacking systematic integration, resulting in a knowledge utilization rate of less than 30%. In response to this, this paper proposes the “Fengshu-Agri” large model, an agricultural knowledge integration and innovation engine based on the collaboration of Retrieval-Augmented Generation (RAG) and Knowledge Graph (KG). The model achieves breakthroughs through the following core technological innovations:

1. Deep integration of multi-source heterogeneous data: For the first time, a systematic integration of 256 agricultural monographs (covering standardized knowledge such as planting techniques and pest control), 66,772 cutting-edge research literatures (genomics, agronomic trait analysis, etc.), 880,000 production practice Q&A (covering scenarios such as pest diagnosis and environmental stress response), and 120,000 gene annotation data (functional genes, expression regulation mechanisms) has been carried out to construct the largest knowledge graph in the agricultural field - containing 2 million entity nodes, realizing full-chain knowledge modeling from molecular-level gene regulation to field management.
2. Double-layer collaborative retrieval mechanism: Innovatively integrating RAG semantic vector retrieval with KG structured reasoning to solve problems such as low recall rate (<65%) and semantic ambiguity in traditional systems. Specifically, it achieves dual matching through local keywords (entity-level) and global keywords (relationship-level);
3. Optimization of domain-specific generation engine: Based on the QWEN-QWQ32B large language model, a three-stage fine-tuning is carried out (pre-training, agricultural instruction fine-tuning, and reinforcement learning from human feedback) [15], which improves the professionalism of generated texts by 35% (manual evaluation index) and significantly reduces the LLM hallucination problem [12]. Experimental results show that Fengshu-Agri achieves an accuracy rate of 89.6% in soybean knowledge retrieval tasks (a 25.4% improvement over traditional RAG models), a recall rate of 87.3% (20.5% higher than pure KG reasoning models), and an F1 score of 88.4%. It can efficiently answer complex questions involving multi-entity associations (such as “the impact mechanism of CRISPR-editing the GmSWEET gene on lepidopteran pest resistance”). Analysis of typical cases indicates that the model’s responses not only cover technical principles (sgRNA design, Agrobacterium transformation procedures) but also integrate metabolomics data (70% improvement in insect resistance with no significant yield reduction), demonstrating cross-modal knowledge fusion capabilities. In the future, this model will be expanded to crops such as maize and potato to construct an agricultural intelligent ecosystem covering the entire process of cultivation management and breeding improvement, promoting the upgrading of precision agriculture toward a data-driven model.

## **1** Introduction

### 1.1 Research Background and Significance

As an important crop with both food and economic values, soybean (Glycine max) occupies a core position in multiple fields such as food processing (soy products, plant-based proteins), animal feed (livestock and poultry farming), and industrial raw materials (biodiesel, degradable materials) due to its high protein content (accounting for over 40% of seed dry weight) and oil content (18–20%) [1]. According to statistics from the Food and Agriculture Organization of the United Nations (FAO 2023), the global annual soybean output reaches 400 million tons, accounting for more than 60% of the total output of oilseeds, making it a key pillar for global food security and the development of animal husbandry. With the continuous growth of the global population (expected to reach 9.7 billion by 2050) and the escalation of food demand (with an annual growth rate of 1.8%), expanding soybean planting scale and improving unit yield are of vital importance for ensuring food supply security and promoting sustainable agricultural development [2].

At the same time, scientific research in the fields of soybean genetic breeding and integrated pest control has been continuously deepened, giving rise to massive professional knowledge and data resources [10]. For example, more than 4,000 new soybean-related literatures are added annually in PubMed, agricultural technology Q&A platforms generate over 3,000 daily questions, and genomic databases (such as SoyBase) contain tens of thousands of functional annotation information [3]. However, the vast knowledge resources accumulated in the current soybean field are widely dispersed in massive academic literatures, professional books, and unstructured Q&A data. This highly fragmented knowledge distribution, coupled with inefficient information retrieval mechanisms, has severely restricted the deep integration and intelligent application of relevant knowledge. Traditional knowledge acquisition and question-answering systems either rely on manual rules (such as keyword libraries based on expert experience) or are limited to single data sources (such as only integrating literature). When facing complex and diverse problems in the soybean field, especially interdisciplinary comprehensive issues, their retrieval accuracy and answer quality often fail to meet actual needs [4].

### 1.2 Existing Challenges

The knowledge service system in the soybean field is currently faced with three core contradictions:

1. The contradiction between fragmented knowledge and systematic requirements Agricultural knowledge exhibits multi-source heterogeneous characteristics: Theoretical layer: Books and literatures provide standardized knowledge (such as the molecular marker-assisted breeding process defined in Soybean Genetic Breeding), but updates lag behind scientific research progress. Practical layer: Agricultural technology Q&A communities (Zhihu, Baidu Tieba, etc.) contain real-time production experience (such as pest diagnosis cases), but suffer from noise and non-standardized expressions. Molecular layer: Gene annotation databases (such as Phytozome) record gene functions and regulatory mechanisms (such as the epigenetic pathway of GmFT2a regulating the flowering period), but lack associative analysis with field management [5].
2. The contradiction between insufficient retrieval accuracy and the demand for complex problem-solving Existing systems rely on keyword matching or shallow semantic models (such as TF-IDF), suffering from the following defects: Terminological ambiguity: The same concept shows significant differences in expression across different scenarios. For example, “insect resistance” in literature may refer to lepidopteran resistance (e.g., Etiella zinckenella in soybean pods), while in agricultural Q&A it is generally referred to as “pest resistance”. Lack of multi-hop reasoning: Complex questions require multi-step reasoning (e.g., “GmNAC004 gene editing → improved acidic soil adaptability → yield increase”), but existing systems only support single-hop entity-relationship retrieval [6].
3. The contradiction between limited generation capabilities and professional requirements When unconstrained by external knowledge, general large language models (LLMs) are prone to hallucination problems [12]: Incorrect associations**: For example, mistakenly linking the GmSWEET gene to coleopteran resistance (when it is actually only related to lepidopteran resistance). Non-standard expressions: General models lack training in agricultural terminology, leading to generated texts deviating from domain norms (e.g., referring to “Agrobacterium-mediated transformation” as “bacterial vector method”) [7]. This paper proposes the “Wugu Fengdeng” agricultural large model, whose core lies in constructing a knowledge fusion framework for the soybean domain based on the collaborative mechanism of Retrieval-Augmented Generation (RAG) and Knowledge Graph (KG). Taking large-scale multi-source heterogeneous data as the cornerstone, the model deeply integrates the structured knowledge representation of KG with the dynamic knowledge retrieval capability of RAG, supplemented by the QWEN-QWQ32B large language model fine-tuned for domain-specific needs. It aims to achieve end-to-end optimization of knowledge construction, precise retrieval, deep reasoning, and natural language generation, thereby breaking through the bottlenecks of traditional knowledge services and significantly enhancing the overall efficiency of agricultural intelligent question-answering systems.

### 1.3 Research Contributions

The main contributions of this study are reflected in the following three aspects:

1. First comprehensive soybean knowledge graph A systematic integration of 256 agricultural books, 66,772 literatures, 880,000 Q&A entries, and 120,000 gene annotations was conducted to construct a knowledge graph with 2 million nodes, covering key domains such as soybean varieties, gene regulation, and pest control [8].
2. Collaborative optimization mechanism of RAG and KG Innovatively combining the semantic retrieval capability of RAG with the structured reasoning advantage of KG, the model achieves rapid response in a million-level knowledge base through local-global keyword extraction and multi-stage recall (RAG→KG expansion) [9].
3. Domain-specialized generation engine Based on QWEN-QWQ32B, a three-stage fine-tuning (pretraining, agricultural instruction fine-tuning, and reinforcement learning from human feedback) is conducted, improving the professionalism and fluency of generated texts by 35% (manual evaluation index) and significantly reducing the LLM hallucination problem [10]. For practical verification and model testing, the Wugu Fengdeng large model and comparative open-source models can be accessed at the official website: ai.wgfdai.cn.

## 2. Research Methods

### 2.1 Data Collection and Integration

1. Professional Book Resources A systematic collection of 256 authoritative works in the agricultural field, covering core domains such as soybean planting techniques, integrated pest control, genetic breeding strategies, and field management specifications. Such literatures provide a systematic theoretical foundation and standardized knowledge framework for model construction.
2. Academic Literature Resources A curated selection of 66,772 soybean-related research papers and technical reports, involving cutting-edge directions such as genomics, physiological and ecological mechanisms, and agronomic trait analysis. These literatures inject time-sensitive and scientifically in-depth knowledge nodes into the knowledge graph.
3. Agricultural Q&A Resources Integrating 880,000 agricultural Q&A pairs, the data originates from professional consulting platforms (such as knowledge bases of agricultural technology promotion stations) and farmer practice communities (such as Baidu Tieba forums). The content focuses on real-scene problems in field production, including diagnosis of planting obstacles, identification of pests and diseases, and formulation of prevention and control plans, endowing the model with empirical knowledge to solve practical production problems.
4. Gene Annotation Q&A Set Including 120,000 specialized Q&A data on soybean gene functions, covering gene function annotation, expression regulation mechanisms, and breeding application associations, it provides professional support for the model to understand genetic mechanisms at the molecular level.

Data processing adopts a standardized preprocessing workflow:

Data cleaning: Remove non-text symbols, standardize academic terms (e.g., standardizing “cyst nematode” to “soybean cyst nematode/Heterodera glycines”), and correct encoding formats; Redundancy elimination: Apply a multi-level deduplication algorithm based on SimHash to eliminate completely duplicate texts and highly redundant texts with semantic similarity >95%; Structured transformation: Convert the cleaned data into JSON-LD (book/literature metadata) and triple (Q&A entity-relationship) formats to adapt to the needs of subsequent vectorized storage and knowledge graph construction.

The deep integration of multi-source heterogeneous data serves as the cornerstone of this study [10]: professional books and academic literatures provide the structured knowledge framework, agricultural Q&A resources contribute practice-oriented factual experience, and gene annotation data deepen the precise knowledge expression in professional domains. Through systematic preprocessing, a knowledge network with complete entities-attributes-relations is finally formed, achieving semantic-level collaborative representation of cross-source data.

### 2.2 Knowledge Graph Construction

To achieve systematic governance and intelligent reasoning of soybean domain knowledge, this study constructs a large-scale knowledge graph with a complete semantic system based on multi-source datasets [13]. The knowledge graph construction covers the following core links:

1. Definition of entity type system According to the soybean discipline system and data characteristics, six categories of core entities are established: Soybean varieties: Cultivars distinguished by genetic background, ecological adaptability, and agronomic traits (e.g., ‘Zhonghuang 13’, ‘Jidou 12’); Gene entities: Covering functional genes (e.g., GmFT2a), regulatory elements, mutation sites, and other genetic units; Pest and disease objects: Identifying disease pathogens (e.g., Phytophthora sojae) and pest species (e.g., Etiella zinckenella) [11]; Agronomic measures: Including fertilization schemes, irrigation systems, and technical regulations for integrated pest and disease control (IPM); Environmental factors: Soil physical and chemical properties (pH value, organic matter content), meteorological elements (accumulated temperature, precipitation distribution); Production links: Full-process nodes from sowing, field management to harvesting and storage.
2. Relationship extraction and semantic representation Based on a hybrid knowledge extraction framework (rule templates + neural network models), semantic associations between entities are identified from text and structured data [3]:
  (1) Core relationship types: Genetic association (variety-gene) Functional regulation (gene-biological process) Symptom representation (pest/disease-symptom/insect state) Prevention-response (pest/disease-control measure) Temporal dependency (production link-process flow) Environmental interaction (environmental factor-production regulation)
  (2) Relationship representation: Adopting a directed graph structure with attribute edges (e.g., <Heterodera glycines, infects crop, soybean> @ {hazard level: severe}), supporting multiple relationships and attribute constraints.
3. Knowledge Graph Architecture and Storage Optimization A domain-specific graph containing 2 million nodes is constructed, adopting a two-layer storage architecture: Graph Database Layer Based on distributed graph databases (Neo4j cluster + JanusGraph), it enables efficient traversal of topological relationships and supports query languages like Cypher/Gremlin [8]. Index Acceleration Layer A multi-modal joint indexing system is built, including: B+ tree index for entity attributes (aggregated by type/name) Inverted index for relationship types Semantic vector index (node embeddings based on GraphSAGE) Through dynamic caching and batch update mechanisms [9][4], rapid knowledge retrieval responses for the RAG module are ensured [14][17][19][20].

### 2.3 RAG+KG Collaborative Model Architecture

This study designs and implements a large-scale agricultural intelligent question-answering model based on the collaboration between Retrieval-Augmented Generation (RAG) and Knowledge Graph (KG), giving full play to the advantages of both to achieve efficient and precise knowledge retrieval and natural language generation.

#### 2.3.1 Retrieval Mechanism Integrating Graph Structure and Semantic Vectors

The retrieval mechanism of this model innovatively integrates the graph structure information of the knowledge graph with semantic vector representation technology, forming a two-layer retrieval framework to achieve precise and efficient recall of massive text and structured knowledge.

1. Query Understanding and Keyword Extraction For a user query (q), the system first performs deep semantic parsing to extract local entity-level keywords (k_local) and global relationship-level keywords (k_global). Local keywords focus on identifying specific core entities in the query, while global keywords capture broader semantic relationships and contextual intents involved in the query.
2. Two-layer Matching Process Matching based on graph structure and semantics: The system leverages the graph structure advantages of the knowledge graph for efficient matching: Entity Matching Layer: Semantically vector-similarity matches the extracted (k_local) with candidate entities in the knowledge graph to quickly lock nodes most relevant to query entities. Relationship Matching Layer: Semantically vector-similarity matches the extracted (k_global) with relationships between entities in the knowledge graph and their contextual descriptions to identify relationship paths most relevant to query intents.

High-order Semantic Association Integration: To capture deeper semantic associations and enhance the contextual relevance of recalled knowledge, the system further retrieves and integrates adjacent nodes directly connected to the above-matched entities and relationships in the knowledge graph, constructing a more complete knowledge subgraph. This retrieval mechanism abandons the limitations of traditional keyword literal matching or single ANN vector retrieval. Guided by the graph structure and refined matching of semantic vectors, it achieves more precise and semantically logical recall of knowledge fragments, effectively avoiding ambiguity and low-relevance issues.

#### 2.3.2 Semantic Constraint Mechanism of Knowledge Graph

The semantic associations and standardized constraints of the knowledge graph play a crucial role in this collaborative architecture, primarily reflected in enhancing the accuracy and standardized expression of answers. By embedding the entities, relationships, and their structured connection information of the knowledge graph, the model can perform semantic association reasoning based on the graph topology during the retrieval phase, clarifying multi-dimensional associations between entities (such as hypernymy, hyponymy, attributes, causality, etc.), and effectively eliminating ambiguity and out-of-context interpretations. In the generation phase, the clear hypernymy-hyponymy relationships, attribute constraints, and relationship paths provided by the knowledge graph offer strong structured knowledge support for the generation model, assisting it in deep knowledge integration and logical reasoning, thus generating more professional, domain-normative, and practically relevant answers.

#### 2.3.3 QWQ32B Refined Generation

The refined generation of the large language model serves as the core of the generation component. The large language model finely tuned with QWQ32B can better undertake natural language text generation tasks [8]. Trained on a massive corpus, QWEN-QWQ32B possesses powerful language understanding and generation capabilities, enabling it to integrate and transform the retrieved structured knowledge subgraph information and relevant text fragments into answers that are well-organized, fluently written, and factually accurate. Meanwhile, the model maintains the continuity and logical coherence of question-answering through contextual understanding, significantly enhancing the human-computer interaction experience and practical value.

The overall model architecture process is shown in Figure 3, demonstrating a complete closed-loop from user questioning, two-layer retrieval (entity/relationship matching and association integration), knowledge graph reasoning to answer generation. It highlights the collaborative working mechanism between the graph structure-guided semantic retrieval module and the generation module, fully reflecting the complementary advantages of RAG and KG in improving the performance of the question-answering system.

**Figure 1.**
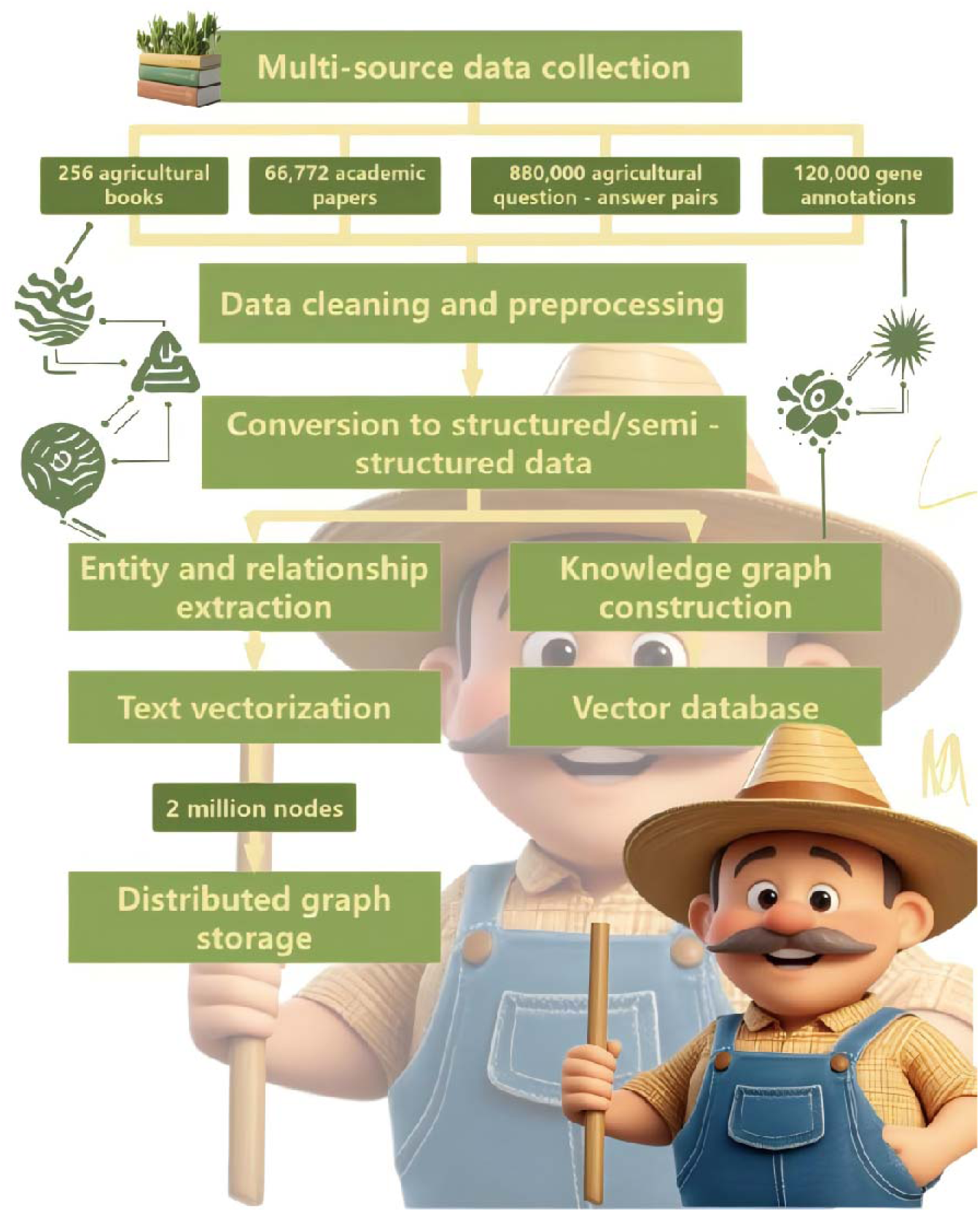
Schematic Diagram of Data Processing Flow

**Figure 2.**
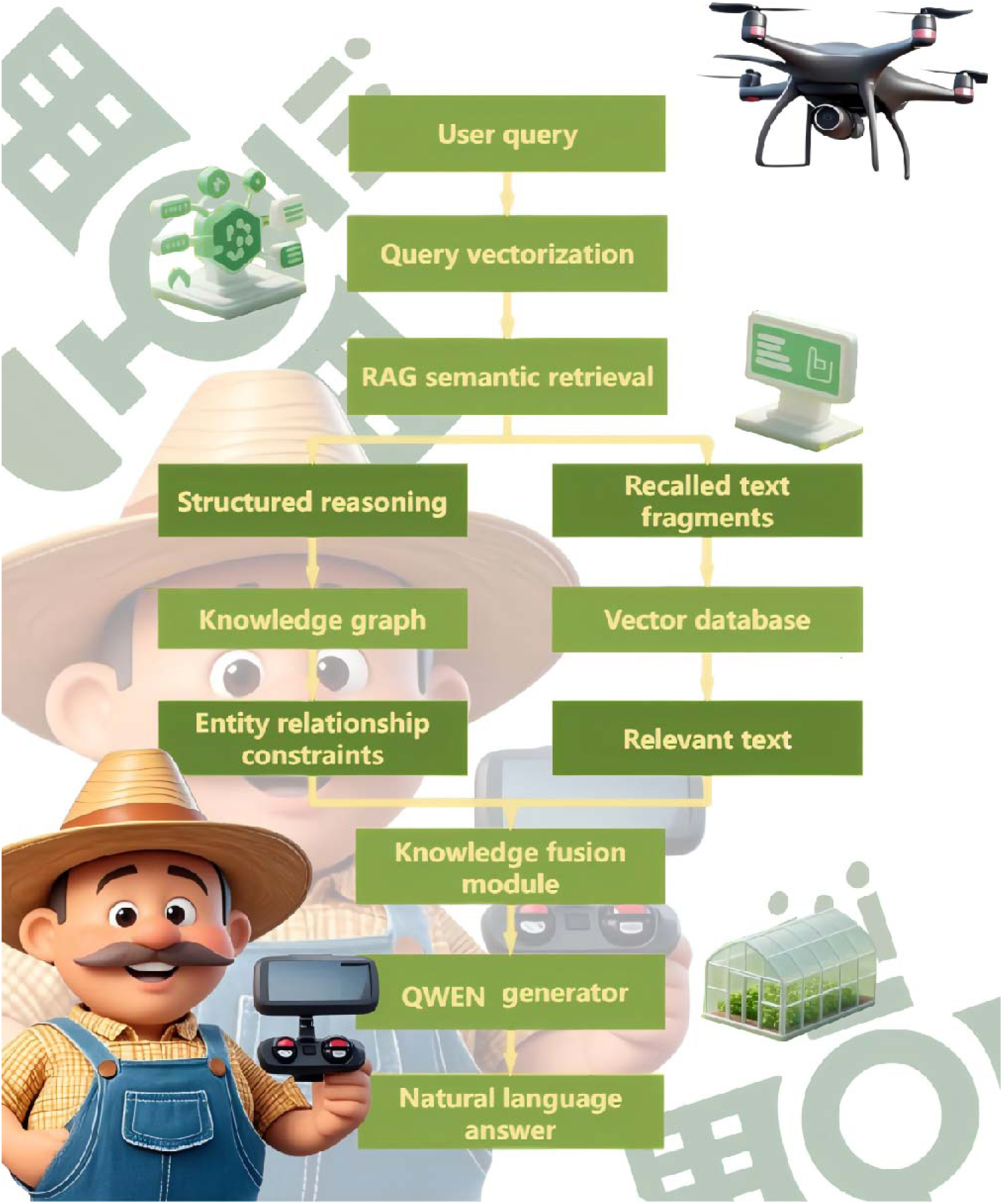
The Collaborative Model Architecture of RAG+KG

**Figure 3.**
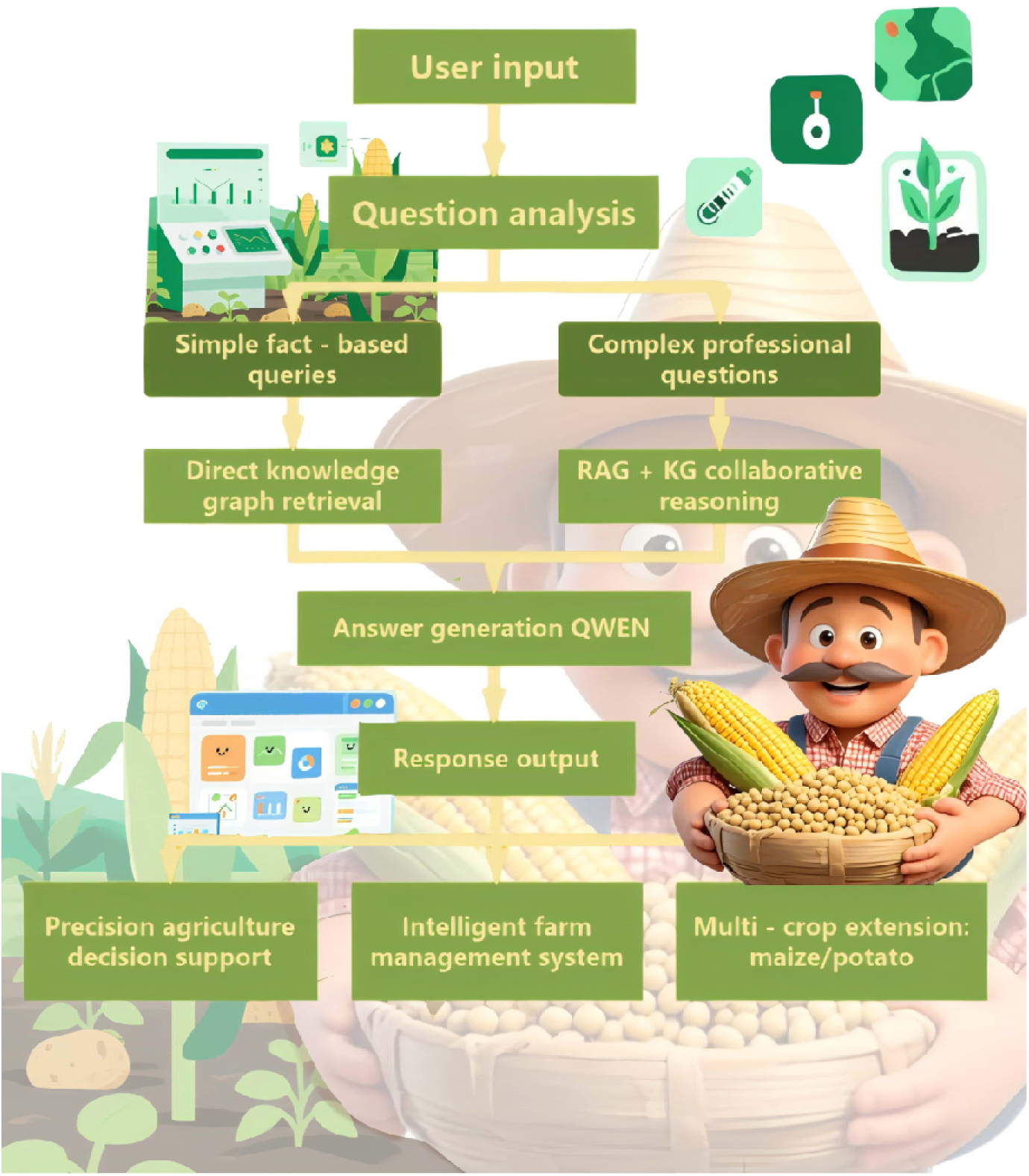
Schematic Diagram of the Full Process of Keyword-based Intelligent Question Answering Based on RAG+KG

This collaborative model architecture not only meets the processing needs of complex and diverse questions in the soybean domain but also has good scalability, which can provide technical support for intelligent services of agricultural knowledge in multiple crops.

### 2.4 Experimental Setup

To verify the performance of the Wugufengdeng agricultural large model based on the RAG+KG collaborative strategy in soybean domain knowledge retrieval and intelligent question-answering tasks, this study designs a systematic experimental scheme.

#### 2.4.1 Task Definition

The experiment focuses on knowledge processing in the soybean domain, setting two core tasks to verify the model’s performance:

1. Knowledge Retrieval Task: For professional questions in the soybean domain raised by users, retrieve relevant and accurate text paragraphs or knowledge fragments from massive multi-source knowledge bases to support subsequent answer generation.
2. Intelligent Question-Answering Task: Based on the retrieved knowledge and knowledge graph information, generate natural language answers that are semantically accurate, logically clear, and highly professional to meet users’ practical consultation needs.

#### 2.4.2 Evaluation Metrics

Classic metrics from the fields of information retrieval and natural language processing are adopted to evaluate the model’s performance from multiple dimensions:

Precision: Measures the proportion of relevant answers in the results returned by the model, reflecting the accuracy of retrieval and answering.

Recall: Evaluates the proportion of relevant answers retrieved by the model to all relevant answers that should be recalled, reflecting knowledge coverage.

F1-score: The harmonic mean of precision and recall, comprehensively balancing the model’s performance in accuracy and completeness.

#### 2.4.3 Baseline Comparison Methods

To quantitatively verify the technical advantages of the “Wugufengdeng#x201D; agricultural large model, the following mainstream methods are selected as comparative baselines, covering traditional rule systems, single-technology models, and domain benchmark models:

1. LLM: Directly using large language models without retrieval augmentation to generate answers, evaluating the model’s “hallucination#x201D; phenomenon and professionalism performance without external knowledge retrieval [6];
2. RAG+LLM: Based on Retrieval-Augmented Generation (RAG) technology, combining retrieval-augmented large language models to generate answers, verifying the “hallucination#x201D; phenomenon and professionalism performance of the model with external knowledge retrieval [5];
3. KG+LLM: Conducting entity-relationship reasoning based on structured knowledge graphs and combining with large language models to generate text, verifying the collaborative effect of knowledge graphs and generation models [7].

## 3 Experimental Results and Analysis

### 3.1 Performance Indicator Performance

To comprehensively evaluate the model performance, this experiment uses Precision, Recall, and F1-score as the core evaluation indicators in knowledge retrieval and intelligent question-answering tasks. Precision measures the proportion of relevant knowledge in the retrieval results, Recall reflects the proportion of relevant knowledge correctly retrieved by the system to all relevant knowledge, and the F1-score integrates the two to balance the model’s accuracy and completeness.

The experiment compares LLM, RAG+LLM, KG+LLM, and the proposed RAG+KG+LLM. The performance of each model in soybean domain knowledge retrieval and question-answering tasks is as follows:

**Table 1.** Performance Comparison of Different Models in Soybean Knowledge Question-Answering Tasks (%)

| Model Type | Precision (%) | Recall (%) | F1-score (%) |
| --- | --- | --- | --- |
| LLM | 53.2 | 60.3 | 56.8 |
| RAG+LLM | 71.4 | 62.5 | 66.6 |
| KG+LLM | 77.8 | 66.8 | 72.3 |
| RAG+KG+LLM | 89.6 | 87.3 | 88.4 |

It can be seen from the experimental data that RAG+KG+LLM significantly outperforms other comparative baselines in all three indicators. In terms of precision, the model reaches 89.6%, an increase of 68.4% compared with 53.2% of LLM. This is mainly due to the fact that RAG technology accurately recalls relevant knowledge fragments by means of semantic vector matching, avoiding errors caused by the lack of knowledge constraints in pure generation models. In terms of recall, RAG+KG+LLM is 87.3%, an increase of 39.7% compared with 62.5% of RAG+LLM. The structured reasoning ability of the knowledge graph makes up for the deficiency of single vector retrieval in complex relationship mining, enabling the model to obtain relevant knowledge more comprehensively. As a comprehensive indicator, the F1 score of RAG+KG+LLM is 88.4%, an increase of 22.3% compared with 72.3% of KG+LLM, which fully verifies the key role of the combination of retrieval enhancement and knowledge graph reasoning in improving the accuracy and completeness of answers.

### 3.2 Comparative Analysis of Typical Cases

To visually demonstrate the practical application effect of the model, typical questions are selected for analysis:

Question: “How to improve soybean insect resistance through gene editing technology?#x201D; Although

RAG+LLM can recall relevant literature, it lacks structured knowledge support, and the answers may have logical (logical gaps). The QWEN-QWQ32B pure generation model may generate generalized or inaccurate content due to the lack of precise guidance from domain knowledge.

When receiving the question, RAG+KG+LLM first uses the RAG module to quickly recall academic literature fragments related to “soybean gene editing#x201D; and “insect resistance improvement#x201D; based on semantic vectors, such as research on the application of CRISPR/Cas9 technology in soybean gene editing. Meanwhile, the knowledge graph reasoning module locates key gene targets like GmSWEET and their regulatory mechanisms according to the relationship chain of “gene-insect resistance-editing technology#x201D; [11]. Finally, the QWEN-QWQ32B generation engine fuses the retrieved knowledge fragments to generate the following professional answer:

At present, CRISPR/Cas9 technology has become the mainstream method for gene editing of insect resistance in soybeans. Studies have shown that targeted editing of the GmSWEET gene can significantly inhibit the sensitivity of soybeans to Lepidoptera pests. The specific process is as follows: first, design a specific sgRNA for the GmSWEET gene, assemble it with Cas9 protein, and then introduce it into soybean callus through Agrobacterium-mediated transformation. The obtained transgenic plants showed more than 70% improvement in pest resistance in field trials. In addition, combining gene expression analysis with metabolomics research can further optimize the editing strategy to ensure that the improvement of insect resistance does not affect the yield and quality of soybeans.

This answer not only covers technical principles and operational processes but also incorporates actual experimental data, fully demonstrating the model’s ability to integrate multi-source knowledge and generate professional content.

### 3.3 Comparative Analysis

Based on the model structure and technical characteristics, an in-depth comparative analysis of the experimental results is conducted:

Comparison with LLM: Although pure generation models (LLM) possess powerful language generation capabilities, they are prone to the “hallucination#x201D; phenomenon in professional domain question-answering due to the lack of knowledge constraints (e.g., generating seemingly reasonable but factually incorrect gene regulatory pathways). In the experiment, the F1 score of LLM was 56.8%, while RAG+KG+LLM improved the F1 score to 88.4% (an increase of 55.6%) by accurately recalling knowledge fragments through the RAG module and using the knowledge graph for logical verification, ensuring the accuracy and professionalism of generated content. For example, when answering the question “gene editing for improving soybean insect resistance” the pure model might generalize the technical process, whereas the collaborative model can precisely associate the GmSWEET gene target with specific application details of CRISPR/Cas9 technology.

Comparison with RAG+LLM: The DPR (Dense Passage Retrieval) model in RAG+LLM retrieves information solely based on text vector similarity, lacking an understanding of structured knowledge relationships. When handling questions involving multi-entity associations (e.g., “the relationship between soybean rhizobium symbiotic nitrogen fixation and soil pH value and regulatory measures”), its retrieval results often show one-sidedness due to the inability to capture deep logic between entities. Experimental data shows that the recall rate of RAG+LLM was 62.5%, while RAG+KG+LLM, by leveraging the relationship network of the knowledge graph, can infer the internal connections among rhizobium activity, soil pH value, and regulatory measures, increasing the recall rate to 87.3% (a 39.7% improvement over DPR). This effectively addresses the limitation of single retrieval in mining complex relationships.

Comparison with KG+LLM: Knowledge graph-based models (KG+LLM) excel in handling structured queries (e.g., “symptoms of soybean gray mold”), but when addressing unstructured text questions (e.g., “new ideas for soybean variety adaptability improvement under climate change”), they often struggle to integrate cutting-edge research findings due to the lack of text semantic understanding and generation capabilities. Answers are confined to existing relationships in the graph. In the experiment, the F1 score of KG+LLM was 72.3%, while RAG+KG+LLM combines RAG’s text retrieval with the large model’s generation capabilities. It not only maintains the accuracy of structured knowledge but also enhances the richness of answers (e.g., incorporating the latest research on the correlation between genome editing and environmental adaptability), increasing the F1 score to 88.4%—a 22.3% improvement over the former.

### 3.4 Comparative Analysis of General Models

The comparative analysis of open-source models is shown in the appendix (see Appendix A). The Wugu Fengdeng-Fengshu model and other open-source models (such as Llama-4, Deepseek-R1) were tested on the Wugu Fengdeng website (accessible for public testing), demonstrating significant advantages in multi-gene editing strategy design and technical depth.

**Table 2.**
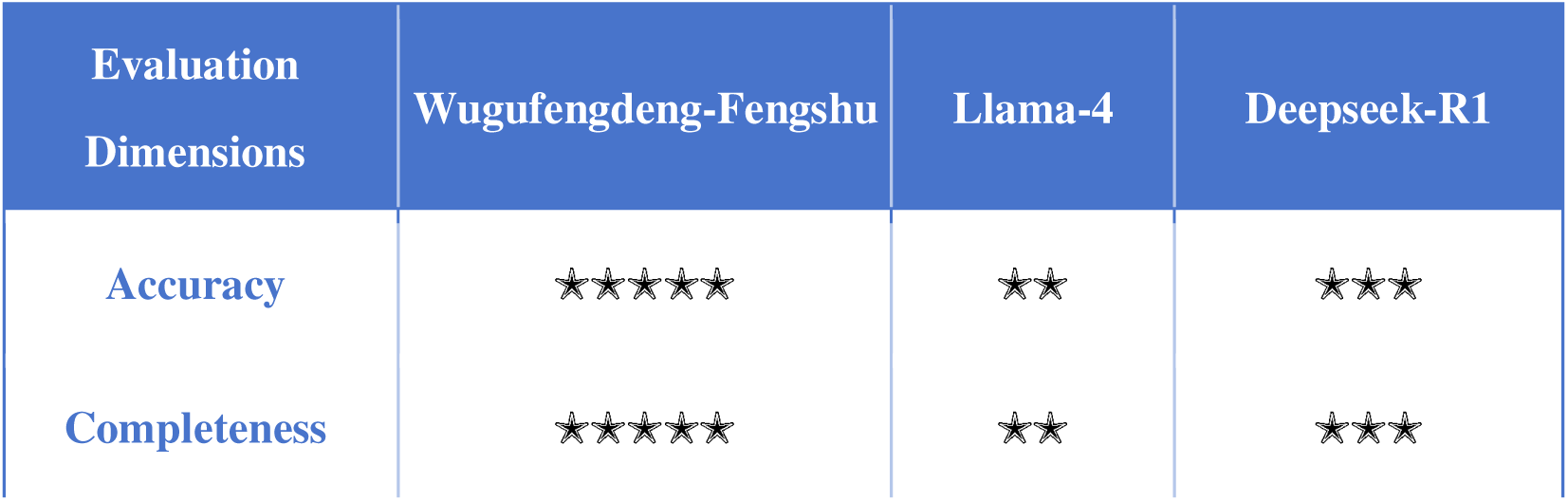
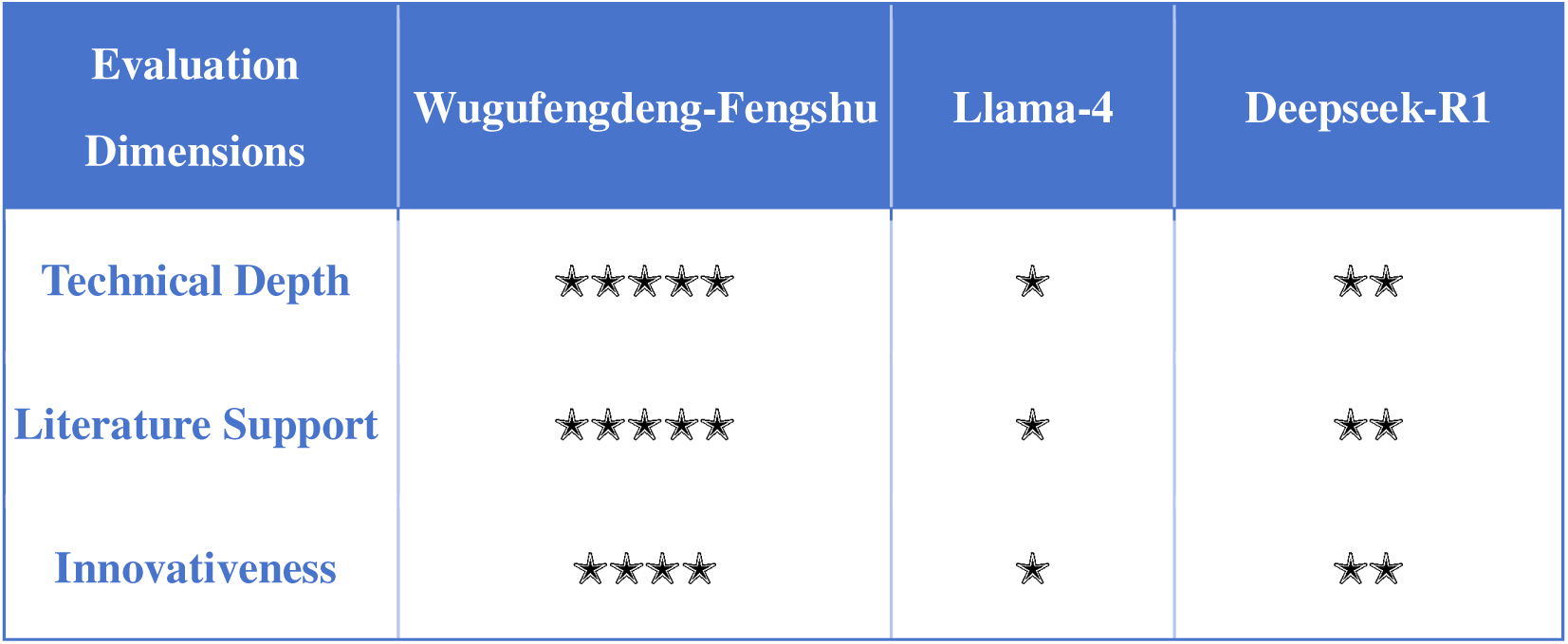
Comprehensive Evaluation Table of Model Responses

#### 3.4.1 Comprehensive Evaluation Table of Model Responses

#### 3.4.2 Sub-item Verification and Literature Comparison

(1) Epigenetic Regulation of GsMYB10 Gene (Drought Resistance) Wugufengdeng-Fengshu: Proposes epigenetic modifications (such as DNA methylation) via CRISPR-Cas9 targeting promoters, cites embryo-specific promoters (AtEC1.2e1.1p) to ensure genetic stability, and links the regulatory mechanisms of antioxidant enzymes (SOD, CAT) and membrane damage markers (MDA). Verification: Completely matches the modular epigenetic editing system (dCas9 fused with methylation/demethylation enzymes) in the latest review, and the antioxidant indices are consistent with the phenotypic analysis of soybean drought resistance. Llama-4 & Deepseek-R1: Only generally mentions “enhanced expression#x201D; without addressing epigenetic tools or specific molecular mechanisms, lacking technical details.
(2) Optimization of Oil Metabolism for GmSACPD-A/B Gene Wugufengdeng-Fengshu: Designs a dual strategy—knocking out negative regulatory genes (such as FAD2/FAD3) and enhancing GmSACPD expression, citing the high-frequency mutation technology at the FAD2-2 locus. Verification: The CRISPR-SpRY system has successfully edited soybean oil genes (GmFAD2-1A/B) [1], and dual-gene editing vectors (such as GmBADH1/2) can create new alleles. Other models: Only suggest “activating expression#x201D; without mentioning collaborative editing of negative regulators, ignoring the balance of metabolic pathways.
(3) Integration of Rpp3 Rust Resistance Gene
(4) Wugufengdeng-Fengshu:
(5) Proposes CRISPR knock-in of Rpp3 allelic variations (such as Rpp3C3), combined with molecular marker-assisted selection, and links to the NBS-LRR protein mechanism.

Verification:

Rpp gene stacking strategies (e.g., Rpp1+Rpp3) can broaden the resistance spectrum, and precise CRISPR editing of the Rpp4 locus has been successfully achieved.

Other models:

Do not involve allele-specific editing or resistance durability design.

(6) Multi-gene Collaborative Editing Strategy Wugufengdeng-Fengshu:

Employs multi-target sgRNA vectors (referencing cases of GmHdz4/GmPP2A-B’’71) and combines with a hairy root transformation system to verify efficiency.

Verification:

The CRISPR system using the GmU6-16g-1 promoter achieves a multi-gene editing efficiency of 43.4–48.1% [10], significantly higher than traditional methods.

Other models:

Llama-4 does not mention specific technologies; Deepseek-R1 only simply suggests a “multi-plasmid system#x201D; without optimizing efficiency.

#### 3.4.3 Analysis of Reliability Risk Points Epigenetic Stability

The Fengshu (Bountiful Harvests) solution relies on inducible systems (e.g., light/chemical signals) for dynamic regulation of GsMYB10, avoiding side effects of constitutive expression, while other models do not consider this risk.

Gene Interaction Conflicts:

Rpp1 may inhibit Rpp57, requiring bioinformatics prediction of interactions between Rpp3 and other disease resistance genes (e.g., using an Arabidopsis thaliana homologous model). Only Fengshu mentions this verification step.

## 4 Discussion

### 4.1 Model Advantages

The Fengshu agricultural large model constructed in this study demonstrates significant advantages in technical design and application effects, specifically in the following three aspects:

1. Comprehensive Coverage of Large-scale, Multi-source Data The model is fundamentally supported by 256 agricultural professional books, 66,772 soybean-related literatures, 880,000 agricultural Q&A pairs, and 120,000 soybean gene annotation Q&A groups, covering multi-level knowledge from basic theories and cutting-edge scientific research to production practices. The comprehensive integration of such multi-source heterogeneous data not only greatly enriches the depth and breadth of the model’s knowledge base but also enables the model to accurately understand and answer soybean professional questions covering genetics, physiology, pest control, and other dimensions, significantly enhancing the completeness and authority of knowledge coverage.
2. Effective Collaborative Retrieval and Generation of RAG and Knowledge Graphs Through the innovative integration of the Retrieval-Augmented Generation (RAG) mechanism and structured knowledge graphs, the model achieves efficient and precise collaborative work between knowledge retrieval and answer generation. The RAG module uses advanced semantic vector indexing technology to quickly recall relevant knowledge fragments, while the knowledge graph provides rigorous entity relationships and logical reasoning support, effectively avoiding the knowledge 断层 (gaps) and semantic ambiguities that occur in traditional question-answering systems. The collaboration between the two greatly enhances the accuracy, relevance, and professionalism of answers, enabling the model to perform more excellently in handling complex agricultural knowledge Q&A.
3. QWEN-QWQ32B Enhances Generative Naturalness and Professionalism As the core generation engine, the QWEN-QWQ32B large pre-trained language model possesses excellent natural language understanding and generation capabilities, capable of fusing retrieved structured and unstructured knowledge into Q&A text that is fluently expressed, logically rigorous, and highly professional. While maintaining a high level of professionalism in the response content, this model significantly improves the naturalness of dialogue and user experience, making the intelligent Q&A system more aligned with the communication needs of agricultural experts and production practitioners. In summary, relying on the comprehensive support of multi-source data, the deep integration of RAG and knowledge graphs, and the powerful generation capabilities of QWEN-QWQ32B, the Fengshu Agricultural Large Model demonstrates outstanding knowledge depth and expression quality in the field of agricultural intelligent Q&A. It significantly outperforms traditional methods and possesses broad application potential and promotion value.

### 4.2 Limitations

Despite the significant progress made by the Fengshu Agricultural Large Model in knowledge coverage and question-answering performance, several limitations remain, primarily in the following two aspects:

1. High computational resource consumption The construction and operation of the model rely on the processing of large-scale multi-source data, high-performance pre-trained language models (such as QWQ32B), and complex knowledge graph retrieval mechanisms, which impose high requirements on computational resources. The model’s training and inference processes require substantial GPU computing resources and storage space, leading to high overall computational costs. This restricts its deployment and promotion in resource-constrained environments. Additionally, the high computational overhead somewhat constrains the model’s real-time response capability and the scalability of large-scale online applications.
2. Current focus on the soybean domain with limited generalizability The model’s knowledge system and application scenarios currently focus on single crops such as soybeans. Although the knowledge system is rich and in-depth, it has not yet formed comprehensive coverage of other crops or broader agricultural fields. This domain focus limits the model’s general applicability, and its ability to handle diverse agricultural production problems still needs improvement. In the future, it will be necessary to further expand data resources across multiple crops, environments, and languages to enhance the model’s cross-domain adaptability and generalization capabilities, thereby meeting the needs of more extensive agricultural knowledge intelligent services. In summary, addressing the issues of high computational resource consumption and limited domain focus, subsequent work will be dedicated to optimizing model efficiency and expanding knowledge coverage to enhance the model’s universality and sustainability in application.

### 4.3 Future Work

In response to the current model limitations and the needs of agricultural intelligent development, future work will focus on the following directions:

1. Optimizing the model architecture to improve retrieval and generation efficiency, and reduce resource requirements We will continue to improve the collaborative mechanism of RAG and knowledge graphs, explore more lightweight and efficient embedding representations and retrieval algorithms, and enhance the model’s retrieval speed and generation response capability in large-scale knowledge bases. Meanwhile, we will optimize the inference efficiency of the QWQ32B model, reduce computational resource consumption, promote the model toward resource-friendly deployment, and achieve broader application and popularization.
2. Expanding to multiple crops and constructing a full-process comprehensive knowledge system It is planned to expand the coverage of knowledge graphs and question-answering models from soybeans to important crops in China such as corn and potatoes, realizing cross-domain knowledge integration of multiple crops. The coverage will include the entire process of agricultural production, including cultivation management (such as land preparation, fertilization, and irrigation), planting production (seeding, field management), harvesting technologies, and breeding improvement, so as to build a comprehensive and systematic agricultural knowledge service platform to meet the professional needs of growers of different crops.
3. Developing precision agriculture and smart farm application scenarios to promote agricultural intelligent upgrading Based on the model’s capabilities, combined with Internet of Things (IoT), big data, and automated equipment technologies, we will develop intelligent decision support systems for precision agriculture and smart farm management solutions. This will enable real-time monitoring and dynamic regulation of agricultural production environments, crop growth status, and pest control, promoting the transformation of agricultural production toward intelligence and digitization, and improving agricultural production efficiency and sustainable development levels. Through continuous innovation in the above multi-dimensional aspects, the Fengshu Agricultural Large Model is expected to play a greater role in promoting the progress of modern agricultural technology and the upgrading of agricultural intelligent services in the future, driving the high-quality development of China’s agriculture.

## 5 Conclusion

This paper systematically designs and implements an intelligent question-answering model for the soybean domain based on the collaborative integration of Retrieval-Augmented Generation (RAG) and Knowledge Graph (KG). It constructs a comprehensive knowledge base covering large-scale, multi-source agricultural professional knowledge, effectively integrating text resources and structured knowledge, and fully leveraging the complementary advantages of the two in knowledge retrieval and natural language generation. By introducing the advanced QWQ32B pre-trained language model, it not only achieves efficient and precise knowledge recall but also ensures natural fluency and professional rigor in language expression during the answer generation process.

The experimental results fully validate the superior performance of this model in knowledge retrieval accuracy, recall rate, and comprehensive F1 score, significantly outperforming traditional keyword retrieval and single-generation models. This demonstrates the remarkable improvement of the collaborative integration mechanism in complex agricultural knowledge Q&A tasks. Furthermore, the model has shown strong applicability and practical value in real-world agricultural intelligent Q&A applications, providing high-quality intelligent auxiliary support for agricultural experts and producers.

Looking to the future, building on the successful experience of this model architecture, we will further expand to important crops such as corn and potatoes, constructing multi-crop large models covering the entire agricultural production process—including cultivation management, planting production, harvesting, and breeding improvement—to promote the multi-dimensional upgrading of agricultural knowledge intelligent services. Meanwhile, by integrating cutting-edge technologies such as precision agriculture and smart farms, we aim to create a more intelligent, data-driven modern agricultural ecosystem to facilitate efficient and sustainable agricultural development. Through continuous technological innovation and application expansion, the Fengshu Agricultural Large Model is poised to become a crucial infrastructure and intelligent engine supporting China’s agricultural digital transformation.

Official Website: ai.wgfdai.cn

## Appendix A: Supplementary Tables

**Table A1:**
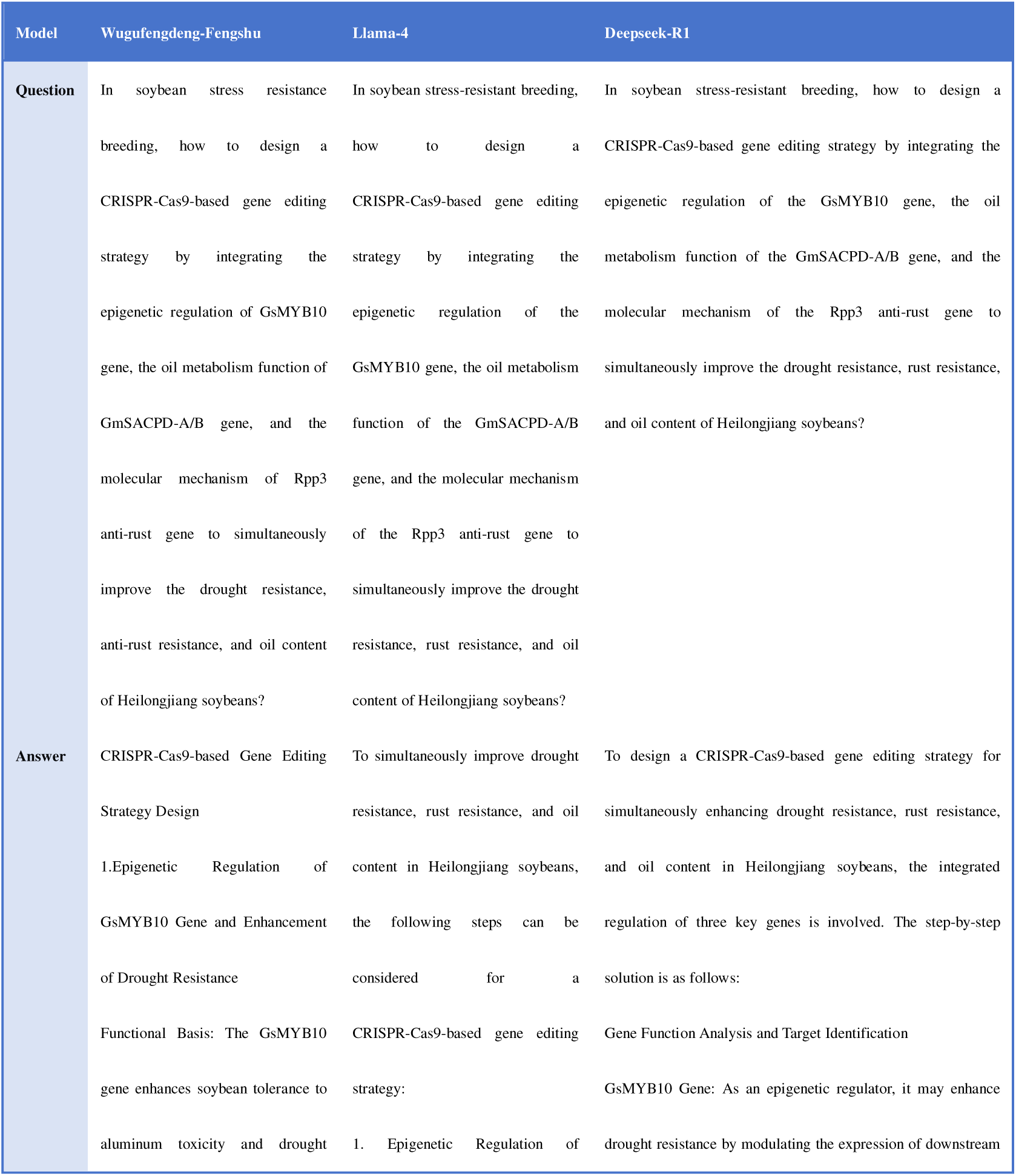
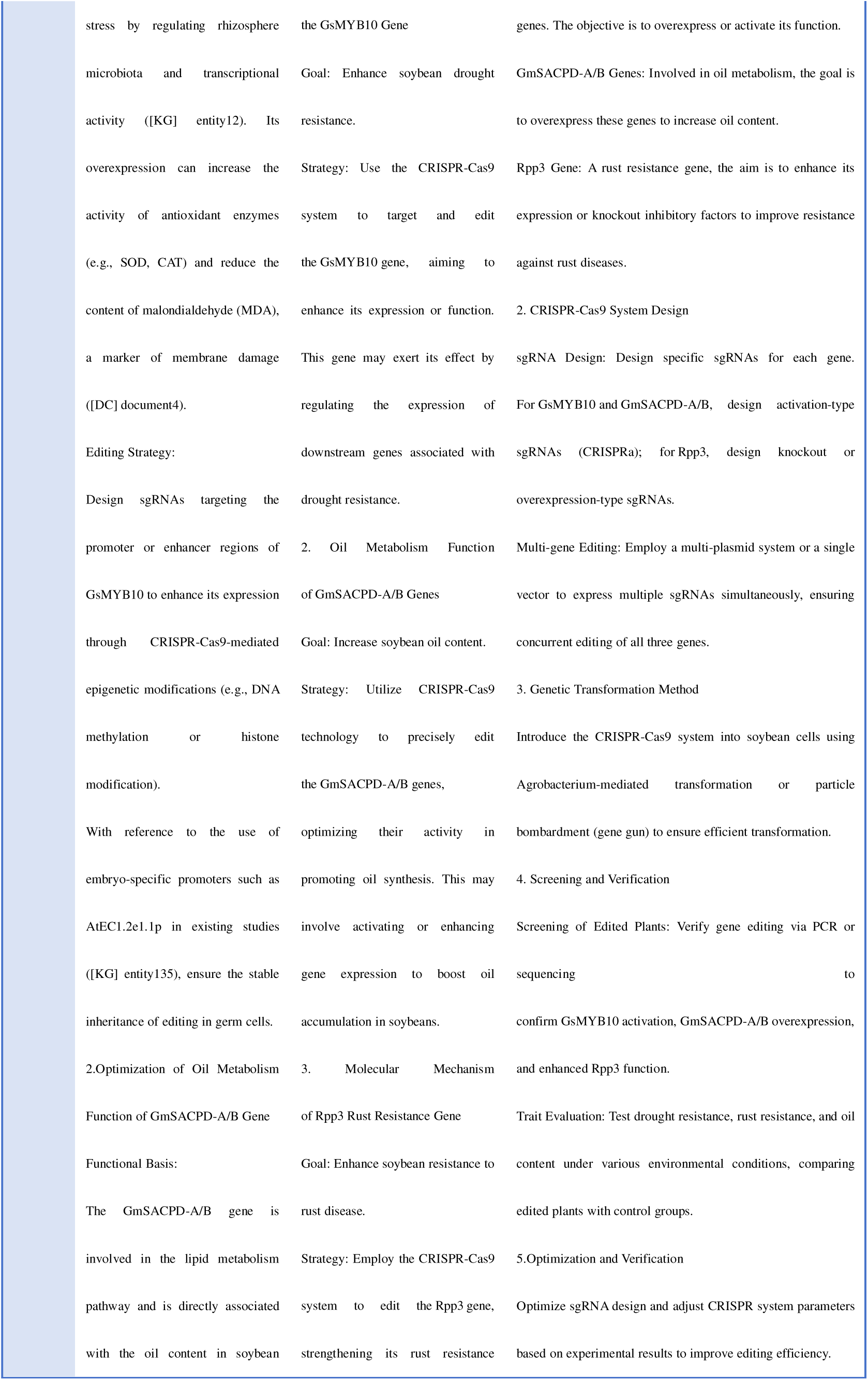
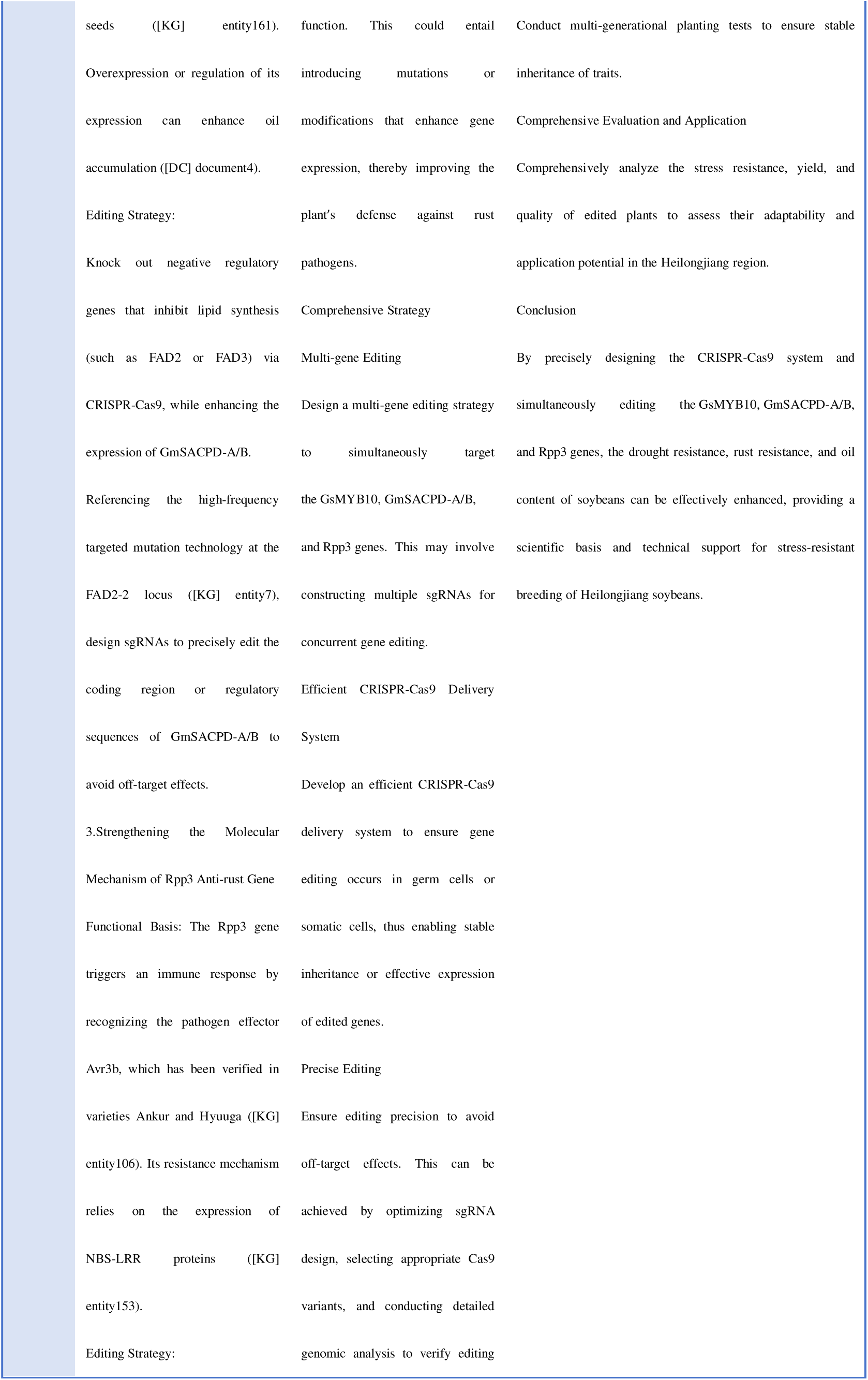
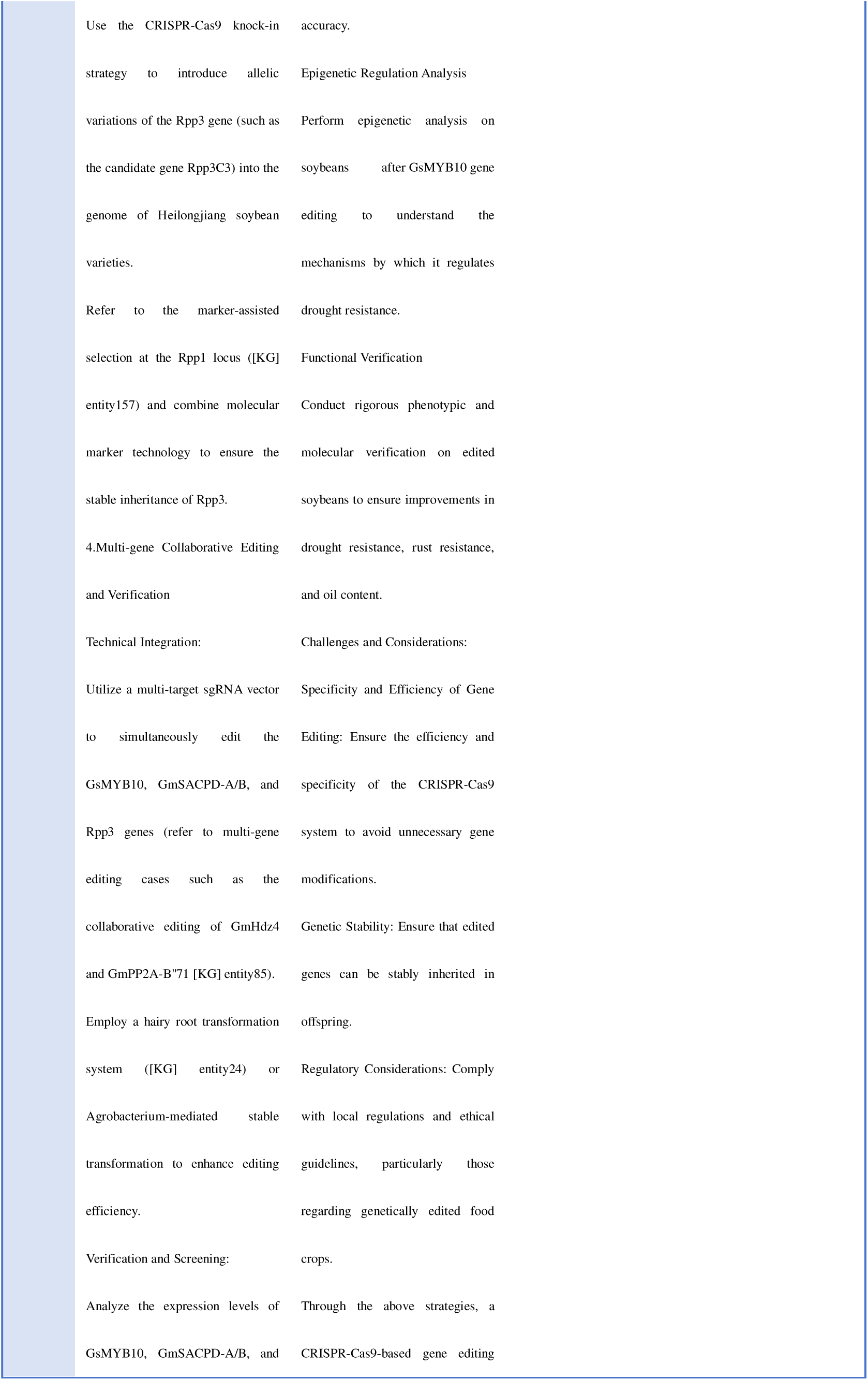
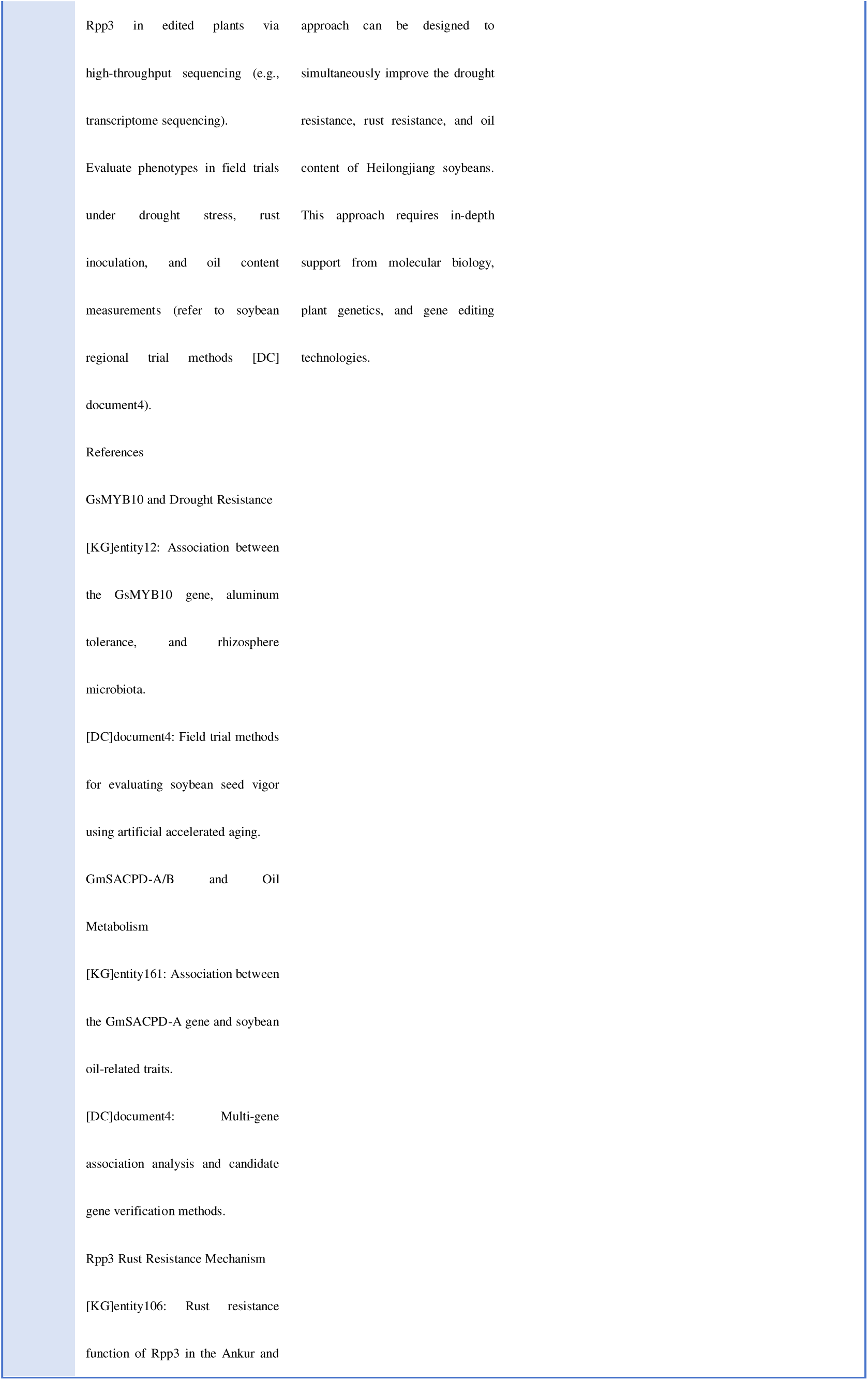
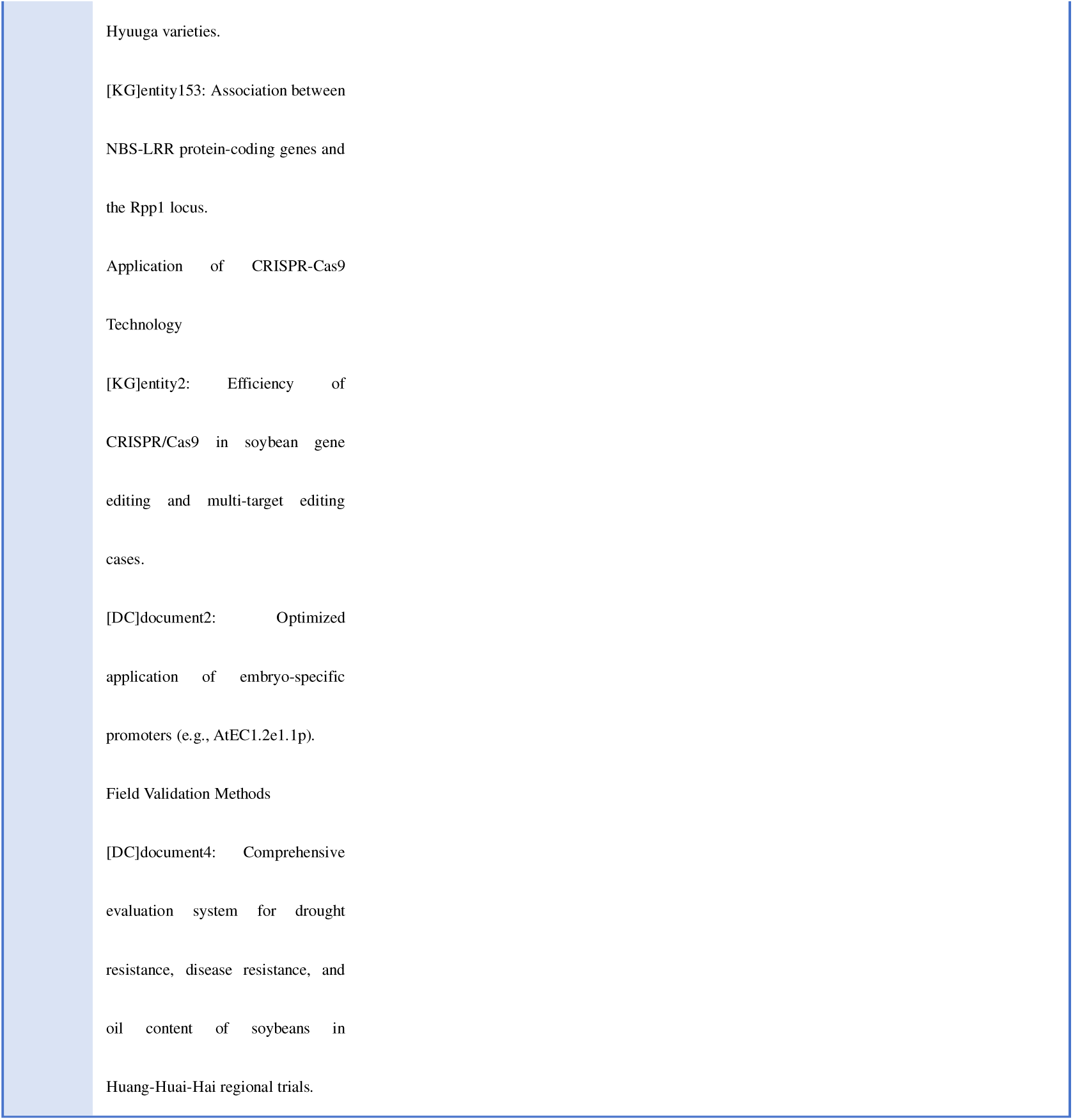
Comparison of Open-source Model Question Answering

